# Unique transcriptional signatures of sleep loss across independently evolved cavefish populations

**DOI:** 10.1101/734673

**Authors:** Suzanne E. McGaugh, Courtney N. Passow, James Brian Jaggard, Bethany A. Stahl, Alex C. Keene

## Abstract

Animals respond to sleep loss with compensatory rebound sleep, and this is thought to be critical for the maintenance of physiological homeostasis. Sleep duration varies dramatically across animal species, but it is not known whether evolutionary differences in sleep duration are associated with differences in sleep homeostasis. The Mexican cavefish, *Astyanax mexicanus*, has emerged as a powerful model for studying the evolution of sleep. While eyed surface populations of *A. mexicanus* sleep approximately eight hours each day, multiple blind cavefish populations have converged on sleep patterns that total as little as two hours each day, providing the opportunity to examine whether the evolution of sleep loss is accompanied by changes in sleep homeostasis. Here, we examine the behavioral and molecular response to sleep deprivation across four independent populations of *A. mexicanus*. Our behavioral analysis indicates that surface fish and all three cavefish populations display robust recovery sleep during the day following nighttime sleep deprivation, suggesting sleep homeostasis remains intact in cavefish. We profiled transcriptome-wide changes associated with sleep deprivation in surface fish and cavefish. While the total number of differentially expressed genes was not greater for the surface population, the surface population exhibited the highest number of uniquely differentially expressed genes than any other population. Strikingly, a majority of the differentially expressed genes are unique to individual cave populations, suggesting unique expression responses are exhibited across independently evolved cavefish populations. Together, these findings suggest sleep homeostasis is intact in cavefish despite a dramatic reduction in overall sleep duration.

## Introduction

Sleep is regulated by homeostatic drive and circadian gating (Archer & Oster, 2015; Borbély, Daan, Wirz-Justice, & Deboer, 2016). Homeostatic sleep pressure increases throughout the day for diurnal animals, while circadian drive aligns periods of rest and activity with light-dark cycles (Archer & Oster, 2015; Borbély et al., 2016). Acute or chronic sleep loss induces recovery sleep, suggesting the presence of a sleep homeostat. While the presence of a rebound in response to sleep deprivation has been described in animals ranging from jellyfish to humans (Anafi, Kayser, & Raizen, 2018; Keene & Duboue, 2018; Libourel et al., 2018; Nath et al., 2017), surprisingly little is known about the molecular basis of this homeostat and even less is known about resiliency of different genotypes to insufficient sleep (Diessler et al., 2018). Further, it is unclear whether the genetic differences that underlie the robust differences in sleep duration are linked to sleep homeostasis or resiliency to insufficient sleep (Keene & Duboue, 2018).

Understanding the functional relationship between sleep duration, homeostasis, and resiliency to insufficient sleep would be aided by investigating the relationship between evolved sleep loss and sleep homeostasis. The Mexican tetra, *Astyanax mexicanus*, has emerged as a leading model for investigating the evolution of sleep (Jaggard et al., 2017; Jaggard et al., 2018; Keene, Yoshizawa, & McGaugh, 2015). *Astyanax mexicanus* are a single species consisting of eyed surface fish that inhabit the rivers of northeast Mexico and Texas and at least 30 populations of conspecific blind, cave-dwelling fish (Bradic et al., 2012; Coghill et al., 2014; Espinasa, Rivas-Manzano, & Pérez, 2001; Herman et al., 2018). Several independent origins of the cave phenotype arose within the past ∼200k years and effective population sizes of independent caves suggest that selection may be an efficient force in shaping cave-derived traits (Bradic et al., 2012; Coghill et al., 2014; Herman et al., 2018). Sleep is reduced compared to surface fish in five cave populations studied to date, suggesting the evolutionary convergence on sleep loss (Duboué, Keene, & Borowsky, 2011; Jaggard et al., 2018; Yoshizawa et al., 2015). Further, distinct neural mechanisms lead to sleep loss between populations, indicating sleep loss evolved independently in different cavefish populations (Jaggard et al., 2017; Jaggard et al., 2018).

Despite this headway in understanding the evolution of sleep duration, the relationship between changes in evolved sleep duration, homeostasis, and cognition and physiological resiliency to insufficient sleep is unclear. Previously, it was reported that one cavefish population displayed a rebound following deprivation and the other tested populations approached significance (Duboué et al., 2011). Given the robust evolutionary changes in sleep duration across populations, it is not clear whether rebound sleep in cavefish is dependent on shared or divergent molecular mechanisms from surface fish. Here, we employ a new method of acute sleep deprivation and find surface fish and three cavefish populations display robust sleep rebound following a single night of deprivation.

A recently sequenced genome in *A. mexicanus* allows for genome-wide analysis of context-dependent changes in gene-expression (McGaugh et al., 2014). While transcriptomic approaches have been used to identify developmentally-specified changes in gene expression in *A. mexicanus* (Gross, Furterer, Carlson, & Stahl, 2013; Hinaux et al., 2013; Stahl & Gross), they have not been applied to identify novel regulators of behavior. The power of comparing between independent populations of cavefish with robust differences in sleep behavior, allows for investigation of mechanisms underlying sleep homeostasis as well as physiological and metabolic responses to sleep deprivation.

Here we compare the behavioral and molecular signatures of sleep deprivation in surface fish and three populations of cavefish. We find that recovery sleep is maintained in cavefish populations despite robust evolutionary loss of sleep duration in undisturbed conditions. Transcriptome analysis revealed unique signatures of sleep deprivation suggesting distinct responses to insufficient sleep across all populations.

## Methods

### Animal Husbandry

Animal husbandry was carried out as previously described (Borowsky, 2008), and all protocols and procedures for this experiment were approved by the IACUC Florida Atlantic University (Protocols A16-04 and A17-32). Fish were housed in the Florida Atlantic University core facilities at 23 ± 1°C constant water temperature throughout rearing for behavior experiments (Borowsky, 2008). Lights were kept on a 14:10 hr light-dark cycle that remained constant throughout the animal’s lifetime. Light intensity was kept between 25–40 Lux for both rearing and behavior experiments. All fry were housed in 250 mL pyrex dishes (0-10 dpf, days post fertilization) or 1L tanks (11-30 dpf) and fed *Artemia* twice daily.

### Sleep deprivation

We conducted the behavioral and RNA-seq experiments with three different cave populations: Pachón, Tinaja, and Molino and one surface population. Surface and cave fry at 30 dpf exhibit differences in sleep duration and architecture (Duboué et al., 2011; Jaggard et al., 2018), thus this is an appropriate stage to assay for variation in sleep rebound. Sleep deprivation was performed by placing fish in 250mL Erlenmeyer flasks on a VWR 2500 multi-vortexer with settings of “4” for speed and “2” for duration. Approximately 20 fry per treatment and population aged 29 dpf were shaken at random intervals < 60 seconds apart since the behavioral definition of sleep is a quiescent period of 60 seconds or more (Duboué, Borowsky, & Keene, 2012; Yoshizawa, 2015). This deprivation paradigm lasted throughout the night hours between ZT14-24 on the night before fish were placed into wells for behavioral recordings the following day. Control animals were held under similar conditions as sleep deprived animals but were not shaken.

### Sleep behavior analysis

sleep deprivation treatment consisted of shaking beakers of fish randomly, at a consistent speed one time per minute throughout a period of 10 hours during the night (ZT 14-24 8pm to 6am). Starting at ZT14, fish in the sleep deprivation treatment were shaken in 200 ml beakers, followed by immediate transfer to 12-well tissue culture plates (BD Biosciences). Control fish were housed under identical conditions but were not shaken. Sleep behavior was recorded in juvenile fish aged 30dpf as previously described (Jaggard et al., 2017; Jaggard et al., 2018). Recording chambers were lit with custom infrared LEDs placed beneath the fish (Jaggard et al., 2019). Videos were recorded using VirtualDub software (v 1.10.4) and were analyzed using Ethovision XT 9.0. All water chemistry was monitored and maintained at standard conditions. Ethovision tracking parameters were set as previously described, and data was processed using custom Perl and Excel scripts (Jaggard et al., 2017; Yoshizawa, 2015). All behavioral recordings were started at ZT0 and recorded for 24 hours. Fish were fed at the start of the behavioral analysis (ZT0). For each treatment and population, 18 fish were analyzed.

### Sample collection for RNA-seq after sleep deprivation

RNA-seq samples were collected from *Astyanax mexicanus* that had been reared in the Keene laboratory at Florida Atlantic University for multiple generations. We conducted the experiment with the same three different cave populations and surface population that were used for the behavioral experiment. Surface parental fish were derived from wild-caught Río Choy stocks originally collected by William Jeffery. Cavefish stocks were Pachón #20, Tinaja 51, and Molino 187.

To minimize non-specific variation, all individuals from each population were collected from the same population-specific clutch (Pachón cave: fertilized from a mating cluster of six parents, Molino cave: from a mating cluster of eight parents, Tinaja cave: from a mating cluster of three parents, Río Choy surface: from a mating cluster of eight parents). Individuals from each population were raised for 27 dpf under standard conditions, (14:10 light-dark cycle), and three days prior to the experiment, fish were transferred into dishes with 12-21 fish per dish in a 14:10 light-dark cycle. Fish were fed ad libitum twice daily in the morning and evening. On days that were non-experiment days, feedings were within a 2-3h window to prevent fish from becoming food-entrained. On experiment days, fish were fed at Zeitgeber time 0 and 12 and were fasted overnight.

As described above, the sleep deprivation treatment consisted of shaking beakers of fish randomly, at a consistent speed one time per minute throughout a period of 10 hours during the night (ZT 14-24, 8pm to 6am). Fish were collected at ZT24 when fish were 30dpf. All individuals were immediately flash frozen in liquid nitrogen upon collection, and stored at −80°C (Figure 1).

**Figure 1.**
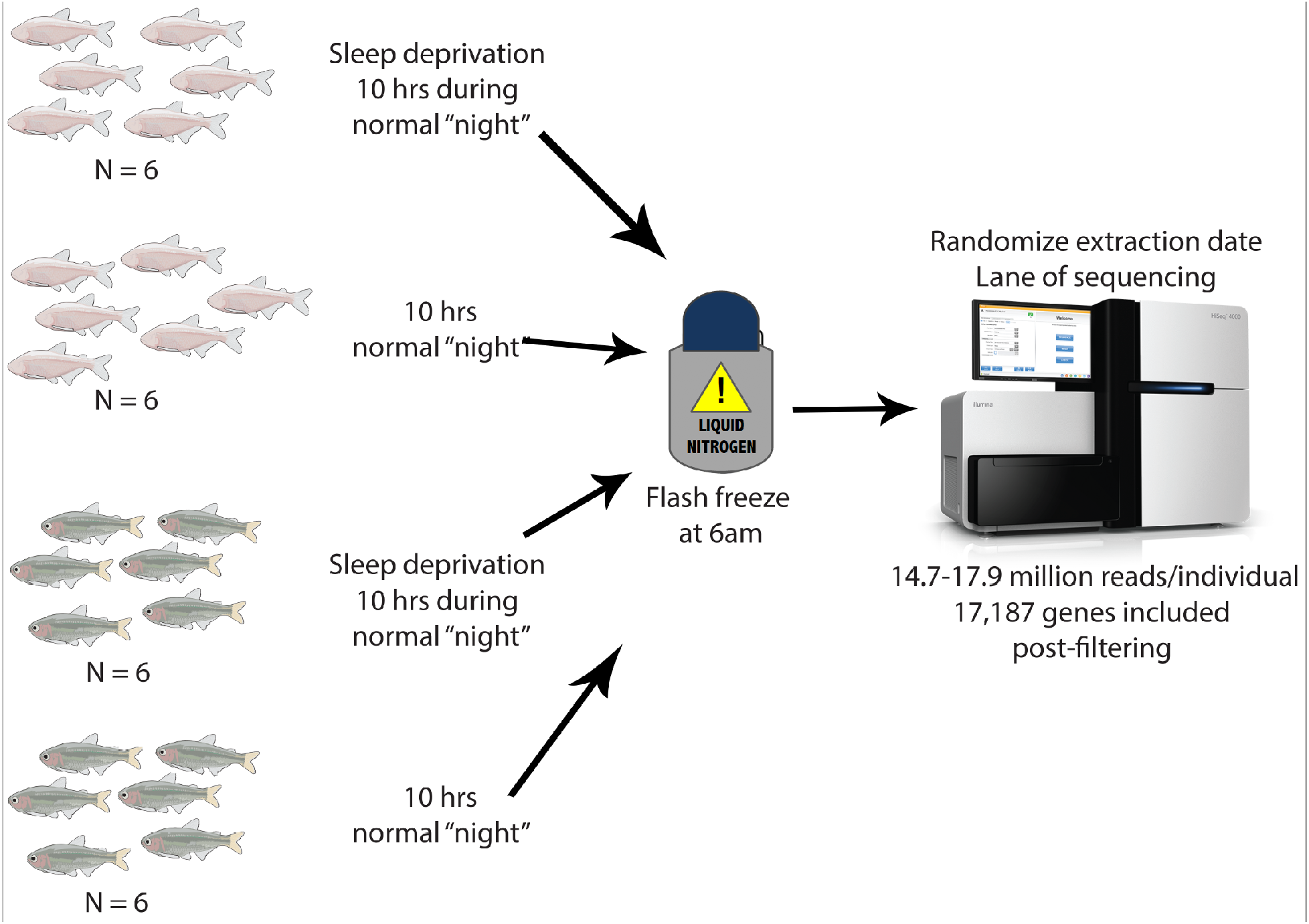
Sampling schematic. We sampled three cave populations: Tinaja, Pachón, and Molino and one surface population at 30 days post fertilization. Fish were sleep deprived for 10hrs during their normal dark period through intermittent shaking.

For the Molino and Pachón populations, control fish were from a spawning of the same parental fish as sleep deprived fish and were exactly 30dpf when sampled, however they were sampled from a clutch that was spawned approximately a week after the sleep deprived treatment fish. We sampled control fish on the same day of as the sleep deprivation fish, but due to a technical error, we could only sequence one control individual from Molino and Pachón from the same sampling day as the sleep deprived fish. We used this single control sample from Molino and Pachón to determine whether collection days drove a substantial amount of variation. MDS plots revealed that there was little difference between these two sampling times (Figure S1).

### RNA extraction, library preparation and sequencing

Protocols for RNA extraction, library preparation, and sequencing were performed as previously described (Passow et al., 2019). To isolate RNA, whole 30dpf fry (∼ 30 mg of tissue) were homogenized and then lysed. Total RNA was extracted using the Qiagen RNAeasy Plus Mini Kit (Qiagen) and then quantified (ng/uL) using NanoDrop Spectrophotometer (Thermo Fisher Scientific), Ribogreen assay (Thermo Fisher Scientific), and Bioanalyzer RNA 6000 Nano assay (Agilent). All individuals were extracted within a week by same researcher. The extraction batch was randomized across populations and treatments (Table S1).

All cDNA libraries were constructed at the University of Minnesota Genomics Center on the same day in the same batch, and 400ng of total RNA was used for mRNA isolation and cDNA synthesis. Strand-specific cDNA libraries were then constructed using TruSeq Nano Stranded RNA kit (Illumina), following manufacturer protocol. Before libraries were pooled for sequencing, library quality was assessed using Agilent DNA 1000 assay on a Bioanalyzer. Only two samples exhibited RIN scores < 9 from the extracted total RNA (Table S1).

To minimize sequencing lane effects, barcoded libraries were pooled with treatment and population spread evenly between lanes. Samples were sequenced across multiple lanes of an Illumina HiSeq 2500 to produce 125-bp paired-end reads at University of Minnesota Genomics Center (Table S1). Little variation was explained by extraction batch or lane of sequencing (Passow et al., 2019). All sequence data were deposited in the short read archive (Table S1).

### Gene expression quantification

The pipelines used to filter and quantify patterns of differential expression were similar to previous published work (Passow et al., 2019), with minor adjustments. In brief, the raw RNA-seq reads were quality checked using Fastqc (Andrews, 2014) and then trimmed to remove adapters and poor quality reads using the program Trimmomatic v0.33 (Bolger, Lohse, & Usadel, 2014). Trimmed reads were then mapped against the *Astyanax mexicanus* Pachón reference genome (version 1.0.2; GenBank Accession Number: GCA_000372685.1; McGaugh et al., 2014) using the splice-aware mapper STAR (Dobin et al., 2013). Stringtie v1.3.3d (Pertea et al., 2016; Pertea et al., 2015) was used to quantify the number of reads mapped to each gene in the reference *A. mexicanus* genome annotation. Finally, to generate the counts matrix, we used the python script provided with Stringtie (prepDE.py; Pertea et al., 2016). Annotations were extracted from the *Astyanax mexicanus* annotation file from Ensembl (Astyanax_mexicanus.AstMex102.91.gtf).

### Variation in gene expression

For quality control, each gene was required to have greater than two counts per million (cpm) and to have counts data for three or more individuals across the sleep deprivation and control samples for all populations. This resulted in 17,187 genes that were analyzed. We identified genes that showed the largest difference in observed gene expression between sleep deprived and standard conditions using the Bioconductor package, EdgeR (Robinson, McCarthy, & Smyth, 2010). We used the calcNormFactors command on our DGEList, and generated multidimensional scaling plots of the top genes using the gene selection method “common.” Given tagwise dispersion and a design matrix, we used glmFIT to fit a negative binomial GLM for each tag and DGEGLM object to perform likelihood ratio test based on the contrast matrix of comparisons. To perform a principal component analysis, we also utilized the log-cpm values of the counts data with the function prcomp(). P-values for differential expression were adjusted based on the Benjamini-Hochberg algorithm, using a default false discovery rate of 0.05 (Love *et al*. 2014). Genes were labeled as differentially expressed if the Benjamini-Hochberg adjusted P-value was less than 0.05 (Tables S2-S5). Log2(Sleep deprived/standard) values were calculated with DESeq2 and exported for further analysis.

### Annotation of differentially expressed genes

We conducted annotation analyses using differentially expressed genes at the 0.05 false discovery rate using PANTHER analysis (Mi *et al*. 2016) (http://pantherdb.org/tools/compareToRefList.jsp). PANTHER analysis was run using only 1:1 orthologs between zebrafish and *Asytanax* with database current as of 2018-12-03. Within the PANTHER suite, we used PANTHER v14.0 overrepresentation tests (i.e., Fisher’s exact tests with FDR multiple test correction) with the Reactome v65, PANTHER proteins, GoSLIM, GO, and PANTHER Pathways. The target list was the zebrafish genes that were 1:1 orthologs to differentially expressed *Astyanax* genes, and the background list was the 1:1 zebrafish orthologs of *Astyanax* genes analyzed for differential expression.

### Selection analyses

We cross-referenced differentially expressed genes to genes likely under selection using the same analysis metrics detailed in (Herman et al., 2018; Yoshizawa et al., 2018). For population genomic measures, we included a core set of whol genome resequenced samples which contained: Pachón cave, N = 10 (9 newly resequenced + the reference reads mapped back to the reference genome); Tinaja cave, N =10; Molino cave, N = 9; Rascon surface, N = 6; and Río Choy surface, N = 9. We required six or more individuals have data for a particular site. For all population genomic measures, we excluded masked repetitive elements, indels (if present in any of the core set of samples), and 10bp surrounding the bases affected by each indel by using the masking_coordinates.gz file available for the *Astyanax mexicanus* genome v1.0.2 though NCBI genomes FTP.

Our procedures for outlier analyses have been described (Yoshizawa et al., 2018), but we briefly recap here. We used hapFLK v1.3 https://forge-dga.jouy.inra.fr/projects/hapflk (Fariello et al., 2013) for genome-wide estimation of the hapFLK statistic of across all 44 *Astyanax mexicanus* samples and two *Astyanax aeneus* samples. HapFLK accounts for hierarchical population structure by building local ancestry trees and detects changes in haplotype frequencies which exceed what is expected for neutral evolution (Fariello et al., 2013). HapFLK outperformed many other statistics (Schlamp et al., 2016) and may be robust to bottlenecks and migration. We augment this measure with metrics of F_ST_, π, and *d*_XY_. We also required that π that was not excessively low in the two surface populations, Rascon and/or Río Choy, (rank of greater than 500 for dense ranking of genes or lowest ∼2.5% of diversity values in the genome) to protect, in part, against calling outliers from inflated relative measures of divergence that were driven by low diversity due to low recombination or sweeps in the surface population. As we are interested in identifying genes under strong selective sweeps in the cave populations, we did not want to exclude genes with low diversity in cave populations. F_ST_ is more sensitive to changes in allele frequency than absolute measures such as *d*_XY_ (Cruickshank & Hahn, 2014).

## Results

To identify expression signatures of sleep deprivation, we performed RNA-seq on surface fish and three populations of cavefish. At 29dpf fish were sleep deprived for 10hrs by mechanical shaking shown to elicit sleep rebound. Fry were sampled for behavioral sleep rebound and RNA-seq at 30dpf.

### Sleep homeostasis behavioral assays

To quantify sleep homeostasis across *A. mexicanus* populations, we sleep deprived fish throughout the night period (ZT14-24) and compared sleep the following day to undisturbed controls (Figure 2A). Surface fish, as well as Pachón, Molino, and Tinaja cavefish displayed a significant increase in daytime sleep following deprivation suggesting sleep homeostasis is intact in all three cavefish populations (Figure 2B, D-G). When sleep measurements were extended into the night following deprivation (ZT14-24), total sleep duration did not differ between control and sleep deprived fish across all four populations tested (Figure 2C, D-G), suggesting the recovery sleep during the day was sufficient to return sleep drive to levels in undisturbed fish. Taken together, these findings reveal sleep homeostasis is intact across three populations of cavefish, despite exhibiting robustly diminished basal sleep levels.

**Figure 2.**
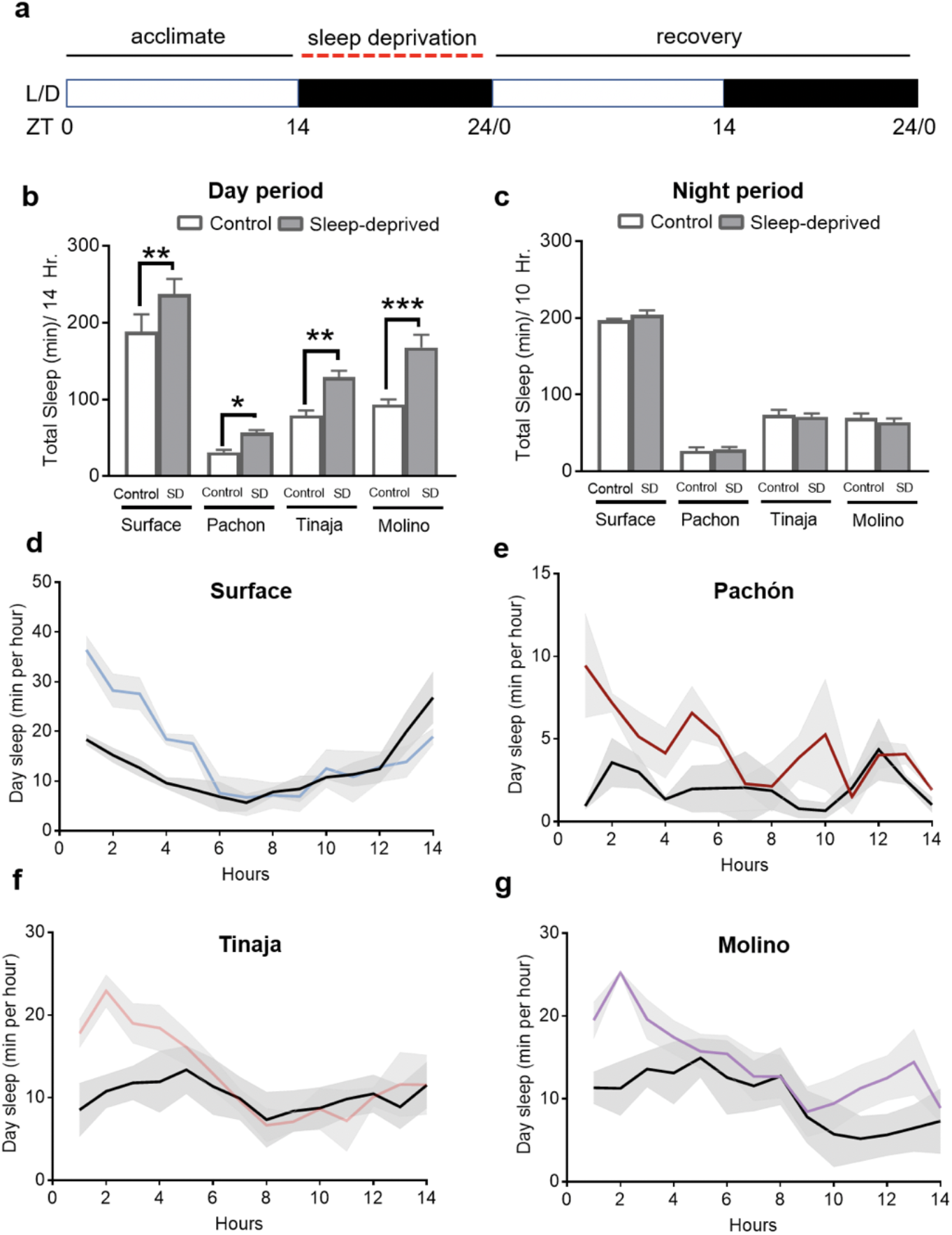
**A)** Following 14hrs of acclimation, fish were sleep deprived for 10 hours (red line) during the night and behavior was measured over the following 24 hrs. **B)** All four groups of sleep deprived fish (grey) slept significantly more than undisturbed controls (white) during the 14hrs of daytime sleep following deprivation due to longer sleep bouts. **C)** No differences in sleep duration were detected between group during the 10 hrs of night following deprivation. Sleep profiles for **D)** surface, **E)** Pachón, **F)** Tinaja, and **G)** Molino over the day following deprivation. Color lines denote sleep derived fish, black lines represent undisturbed controls. Grey represents standard error of the mean.

### Mapping statistics

On average we sequenced 16.72 million reads per sample (median = 16.77 million, min = 13.36, max = 19.88) from whole, individuals at 30 dpf (Table 1, Table S1, Supplemental materials). Pachón cavefish exhibited slightly higher percentage of reads mapped compared to all other populations, as would be expected given reference sequence bias. Total yield of reads were not significantly different between treatments (Wilcoxon test W = 252.5, p-value = 0.4705). On average, 88.96% reads mapped per sample to the *Astyanax mexicanus* genome (median = 88.9%, range 85.06% - 91.59%). Filtering of the gene counts matrix to include only transcripts with greater than two cpm and three or more individuals with counts data resulted in 17,187 genes used for differential expression analysis and clustering analysis, except where otherwise noted. Libraries were normalized by size using calcNormFactors (Figure S2), and samples did not exhibit clustering due to lane of sequencing or extraction batch (Figure S3, S4).

**Table 1.** Summary statistics of samples and sequencing. Standard deviation follows the mean number of raw reads (in millions). Upregulated and downregulated are given as population specific/total differentially expressed genes after correction for multiple testing. The percent that follows pertains to population specific genes.

|  |  |  | Mean # |  |  |
| --- | --- | --- | --- | --- | --- |
|  | Population - Treatment | N | Raw Reads | upregulated | downregulated |
|  | Surface - sleep deprived | 6 | 16.90±0.52 | 124/143 (86.7%) | 101/121 (83.5%) |
|  | Surface - control | 6 | 17.18±1.84 |  |  |
|  | Molino cave - sleep deprived | 6 | 17.91±1.07 | 66/91 (72.5%) | 111/149 (74.5%) |
|  | Molino cave - control | 6 | 16.48±1.75 |  |  |
|  | Pachón cave - sleep deprived | 6 | 17.45±1.45 | 45/66 (68.2%) | 54/81 (66.17) |
|  | Pachón cave - control | 6 | 17.31±1.06 |  |  |
|  | Tinaja cave - sleep deprived | 6 | 14.76±0.60 | 188/230 (81.7%) | 106/137 (77.4%) |
|  | Tinaja cave - control | 6 | 15.75±1.26 |  |  |

### General patterns in expression response to sleep deprivation

After correcting for multiple testing, the number of significantly differentially expressed genes between sleep deprived and control groups is < 2.1% of the 17,187 genes used for differential expression analyses in all populations (Table 1). Multidimensional scaling analysis showed that the major axis of differentiation among the top 500 genes was ecotype (Figure 3) and explained 33% of the variation, suggesting that expression profiles of cave populations are more similar to each other than to surface fish. The second principal component accounted for 12% of the variation in the data and separated the caves into two clusters containing 1) Pachón-Tinaja and 2) Molino, which represent two separate lineages of cavefish (Herman et al., 2018). Thus, ecotype and population (and not sleep deprivation treatment) drove the majority of variation in the complete dataset. Since population is such a strong driver of expression, to isolate the effect of sleep deprivation on expression profiles, we separated the populations into individual MDS plots (Figure S5, S6, S7, S8).

**Figure 3.**
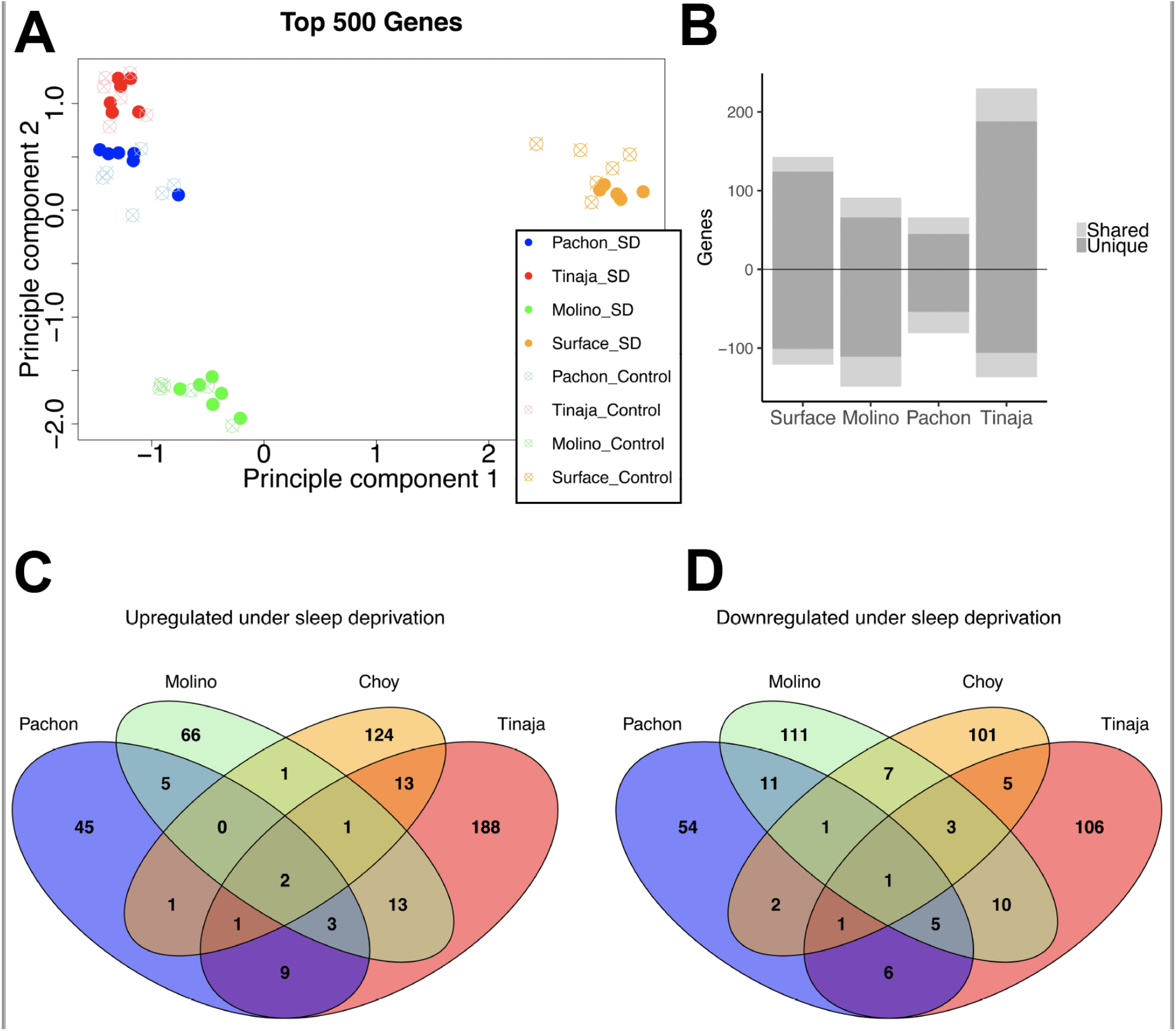
**A)** MDS plot of 500 Top genes between control and sleep deprivation treatments the four populations normalized for library size. PC1 clearly demarcates population of origin. **B)** Number of genes differentially expressed between control and sleep deprived treatments. Above zero is upregulated in sleep deprived samples and below zero is downregulated in sleep deprived samples. Light grey = shared with at least one other population, Dark grey = not shared among other populations. Overall few genes are differentially expressed in response to sleep deprivation and differentially expressed genes are not shared among populations. **C)** Venn-diagram of overlap in differentially upregulated genes in the sleep deprivation treatment between all four populations. **D)** Venn-diagram of overlap in differentially downregulated genes in the sleep deprivation treatment across all four populations.

Across population-specific MDS plots, sleep deprivation treatment accounted for 12-16% of the variation in the dataset. However, individual populations exhibited different signatures. Most notably, Pachón cave population exhibits little differentiation between sleep deprived and control samples (Figure S5) compared to the other populations (Figure S6, S7, S8).

### Divergent responses across cave and surface populations in response to sleep deprivation

Despite surface fish exhibiting greater than 3-fold longer total sleep than cavefish at this stage of development (Duboué et al., 2011 and Figure 1), the 10-hour sleep deprivation treatment did not result in a significant increase in the total number of differentially expressed genes in surface fish compared to any of the cave populations. Surface fish exhibited ∼1.6 and ∼2.2 fold the number of upregulated genes in response to sleep deprivation than Molino and Pachón cavefish, respectively (Table 1). Surface fish also experienced ∼1.5 fold the number of downregulated genes than Pachón cavefish (Table 1). However, Tinaja cavefish exhibited a greater total number of upregulated genes than the three other populations (Table 1). Thus, we did not observe a direct pattern that evolved sleep loss in cave populations results in fewer differentially expressed genes in response to sleep deprivation, fortifying the behavioral findings that sleep homeostasis is largely intact in cavefish population.

While surface fish did not exhibit more exacerbated transcriptional responses to sleep deprivation than cavefish, the surface population exhibited the highest number of uniquely differentially expressed genes than any other population (83.5-86.7%, Table 1), suggesting surface-specific responses to sleep deprivation. More cave-specific overlap in differentially expressed genes, though, could be because the analyses included more cave populations than surface populations.

### Population-specific changes in response to sleep deprivation

Across all cave and surface populations, more differentially expressed genes were found to be shared than estimated by chance, suggesting a degree of conserved responses to sleep deprivation. Between cave populations, the number of concordant upregulated and downregulated genes was significantly greater than random for all comparisons (p ≪ 0.0001; Figure 3). Between cave and surface comparisons, there was significant overlap, but to a lesser degree than in comparisons considering only cave populations (p ≤ 0.007, for all cave-surface comparisons). As would be expected, the number of genes showing discordant patterns (upregulated in one population and downregulated in another) was not significantly different from random for all pairwise-population comparisons (p > 0.145 in all cases). Therefore, across populations more genes were overlapping in concordant transcriptional response than expected by chance alone.

While the overlap was significantly more than expected by chance, few differentially expressed genes were shared between populations. For example, between 5.0-7.8% of differentially expressed genes overlapped in the same direction among pairwise comparisons of caves, and 1.4-2.4% of differentially expressed genes overlapped across all three cave populations. While we found significant overlap in expression responses between populations, 67-82% of differentially expressed genes are unique to specific cave populations. This lends support to the hypothesis that response to sleep deprivation and sleep homeostatic processes may exhibit strong intraspecies differences among caves.

### Functional enrichment analysis

Very few functional categories were significantly enriched after correction for false discovery rate across all populations (FDR; Tables S6-S13). For cave populations, gene ontology analyses indicated that upregulated genes were significantly enriched for oligopeptide transmembrane transport activity (Pachón) and intermediate filaments cytoskeleton organization associated with wound healing (Tinaja). One of the consistently upregulated genes in past mammalian studies of sleep homeostasis is *activity-regulated cytoskeletal associated protein* (*arc*) (Diessler et al., 2018). While this gene is not annotated in any fish genome on Ensembl, it is notable that cytoskeleton-related processes are enriched in genes upregulated in response to sleep deprivation in Tinaja cavefish. Downregulated genes for cave populations were associated with winged helix/forkhead transcription factors in Molino (p = 0.054, after FDR correction) and nitrogen compound transport in Tinaja. In the surface population, upregulated genes are significantly enriched for transmembrane transport functions of a variety of molecules, but downregulated genes exhibit no significant functional enrichment. The different GO categories for each population support the notion that sleep deprivation evokes variable intraspecific physiological responses across different *A. mexicanus* populations.

### Selection analyses

Several differentially expressed genes also exhibit signatures of positive selection (Table S14). Notably, *aif1l* (*allograft inflammatory factor 1 like*) is downregulated in Molino and Pachón cavefish in response to sleep deprivation and exhibited population genetic parameters suggestive of a recent selective sweep in both populations (e.g., significant with HapFLK analyses, in the top 5% of F_ST_ values across the genome between surface and Molino populations, in the top 10% of F_ST_ values across the genome between surface and Pachón populations, and one of the least diverse genes across the genome in both cave populations). Knockout mice of *aif1l* exhibited deformities in endocrine/exocrine, hematopoietic, immune, integument, and liver/biliary systems (Dickinson et al., 2016), suggesting a pleiotropic link between extended wakefulness and liver and immune function in cavefish. Next, *slc17a8* is downregulated in Tinaja cavefish in response to sleep deprivation and is involved in equilibrioception, neuromast hair cells, neuron-neuron synaptic transmission, posterior lateral line neuromast hair cell development, startle response, and vestibular reflex (Obholzer et al., 2008). Population genetic parameters of *slc17a8* suggest a recent selective sweep in Tinaja (significant with HapFLK, in the top 5% F_ST_ outliers between Tinaja cave and Rascon surface, while being among the lowest diversity genes in Tinaja). The down regulation of lateral line components in response to sleep deprivation is especially interesting since sleep duration in at least one cavefish population is restored with the ablation of the lateral line (Jaggard et al., 2017). Our results suggest that dampening signals from the lateral line may be a response to sleep pressure. Interestingly, Pachón cavefish exhibited significant upregulation of the retinal photoreceptor-associated gene *phosphodiesterase 6G* (*pde6gb*) which is highly enriched in the pineal gland relative to the brain of larval and adult zebrafish (Toyama et al., 2009), suggesting *pde6gb* may play a role in circadian physiology. This gene exhibited metrics of a recent selective sweep in Pachón (e.g. significant with HapFLK analysis, in the largest 20% Dxy and largest 5% Fst for Pachón-surface comparisons and exhibits very low diversity in Pachón). The three cases highlighted here suggest that selection in cave populations may have shaped liver, lateral line, and photoreceptor functioning, and these traits pleiotropically impacted sleep phenotypes or vice versa.

## Discussion

While sleep duration is widely studied across animal species, sleep homeostasis and baseline sleep phenotypes are likely functionally distinct (Anafi et al., 2018; Joiner, 2016; Keene & Duboue, 2018). A central question is whether the truncated sleep duration that has evolved in cavefish (Duboué et al., 2011; Jaggard et al., 2018) also impacted sleep homeostasis or whether these processes are independently regulated. Here we implemented an acute sleep deprivation treatment to examine the behavioral and molecular effects in surface fish and three independently-evolved cavefish populations. Our behavioral analysis confirms previous findings that sleep homeostasis is intact across cavefish populations (Duboué et al., 2011), supporting the notion that baseline sleep phenotypes are regulated by separate processes from sleep homeostasis (Allada, Cirelli, & Sehgal, 2017). Sleep significantly deprivation impacted expression of only a few hundred genes across populations, and different cavefish populations and surface fish exhibit discordant gene expression responses to the same sleep deprivation treatment. Thus, our results demonstrate intraspecific variation in molecular mechanisms of sleep pressure in response to sleep deprivation.

Sleep deprivation is an effective way to induce sleep rebound and results in a host of pathophysiological and cognitive deficits (Cappuccio, D’elia, Strazzullo, & Miller, 2010; Cedernaes et al., 2018; Leproult, Holmbäck, & Van Cauter, 2014). Our analysis comparing gene expression in control and sleep deprived fish revealed differential gene expression in only a few hundred genes in response to sleep deprivation across all populations. However, we observed three genes that were differentially expressed across all cave and surface populations, indicating they are part of a generalized response to sleep deprivation: *heat shock protein alpha-crystallin-related, b11* (*hspb11*) and *neutral cholesterol ester hydrolase 1a* (*nceh1a*) were upregulated and transcription factor *E74-like factor 3* (*elf3*) was downregulated in response to sleep deprivation. Heat shock proteins and other molecular chaperones are often upregulated after sleep deprivation as an indicator of cellular stress, suggesting this is an evolutionarily conserved response to sleep deprivation (Allada et al., 2017; Cedernaes et al., 2018; Mackiewicz et al., 2009; Uyhelji et al., 2018). *Neutral cholesterol ester hydrolase 1a* (*nceh1a*) promotes adipogenesis (Homan, Kim, Cardia, & Saghatelian, 2011), and is strongly implicated in atherosclerotic lesions (Igarashi et al., 2010; Okazaki et al., 2008), indicating that *nceh1a* may be a candidate for mediating the well-documented relationships between short sleep duration and advancement of atherosclerosis and weight gain (Cedernaes et al., 2018; Davies et al., 2014; Levy et al., 2012; Nakazaki et al., 2012). *E74-like factor 3* (*elf3*) is the only gene significantly downregulated in response to sleep deprivation in all populations. Interestingly, this gene is involved in inflammatory response (Conde et al., 2016), epithelial cell differentiation (Kwon et al., 2009), and is a cancer gatekeeper (Gajulapalli et al., 2016; Shatnawi et al., 2014; Yachida et al., 2016; Yeung et al., 2017), suggesting a molecular link between sleep disruption and increased cancer risk (Blask, 2009; Haus & Smolensky, 2013). Together, these genes may be conserved factors that are regulated in accord with sleep drive and deprivation across *A. mexicanus* populations.

All genes that were not differentially expressed in the surface population, but were significantly upregulated across all cave populations were linked to sleep or circadian function. *Carnitine palmitoyltransferase 1B* (*cpt1b*) variants are associated with narcolepsy in humans and upregulated *cpt1b* may increase hypocretin activity (Han et al., 2012; Miyagawa et al., 2008), consistent with previous work demonstrating that increased hypocretin is highly associated with short sleep duration in cavefish (Jaggard et al., 2017; Jaggard et al., 2018). *aarF domain containing kinase 5* (*adck5*), was upregulated across all cave populations in response to sleep deprivation and human variants are associated with insomnia symptoms (Lane et al., 2017). *Telethonin* (*tcap*), a component of cardiac sarcomeres, is regulated by circadian components CLOCK and BMAL1 (Podobed, Alibhai, Chow, & Martino, 2014), supporting the close relationship between sleep homeostasis and circadian cycles suggested by other work (Allada et al., 2017; Borbély et al., 2016). In contrast, none of the genes that were significantly downregulated in response to sleep deprivation across all caves have direct links to sleep or circadian function (e.g. *gamma-glutamyl hydrolase-like, ETS homologous factor*, or *cathepsin K*), though *transmembrane and coiled-coil domain family 1* (*TMCC1*) may potentially be involved in endoplasmic reticulum stress or downregulation of protein synthesis, which are common responses to sleep deprivation (Mackiewicz et al., 2009) (Zhang et al., 2014).

After sleep deprivation, five genes were upregulated in surface fish while being downregulated in at least one cave population. In contrast, no genes were downregulated in surface fish while being upregulated in any cavefish population, lending support for the idea that cavefish may be more resilient to extended wakefulness than surface fish. All genes downregulated in Molino cavefish and upregulated in surface fish play critical roles in glucose homeostasis. In mammals, increased expression of *sodium-coupled neutral amino acid transporter 3* (*slc38a5*) triggers pancreatic alpha cell proliferation (Kim et al., 2017) which secrete glucagon to elevate the glucose levels in the blood. In response to sleep deprivation, surface fish upregulate *slc38a5a*, which is typically upregulated when glucose levels drop and circulating amino acid levels increase. The ultimate effect of upregulating *slc38a5a* in surface fish is an expected increase in glucagon and subsequently, elevated circulating glucose levels. In contrast to surface fish, Molino cavefish downregulate *slc38a5a* in response to sleep deprivation which would ultimately lead to a decreased in glucagon and subsequently, decreased circulating glucose levels. Next, *proteinase-activated receptor 2-like* (*par2b* aka *f2rl1.2*) is involved in a variety phenotypes, but recent evidence suggests that downregulation of *par2b*, as seen in Molino cavefish, would result in reduction of generation of glucose from non-carbohydrate sources (e.g. gluconeogenesis), ultimately leading to a reduction in circulating glucose levels (Wang et al., 2015). Lastly, knockout *CCAAT/enhancer-binding protein beta-like* (*cebpb*) mice exhibit reduced hepatic glucose production through glycogenolysis (Liu et al., 1999), suggesting that reduced expression in Molino cavefish may result in the reduction of the conversion of glucagon to glucose. Similar to surface fish, *cebpb* is upregulated in response to sleep deprivation in rats (Elliott et al., 2014) (Cirelli, Faraguna, & Tononi, 2006) suggesting increased glucose production. These expression differences are notable as cavefish and surface fish experience different physiological responses to starvation (Jaggard et al., 2017). Surface fish implement sleep deprivation upon starvation, presumably to increase time for food searching, while Molino and Pachón cavefish increase sleep upon starvation, presumably to conserve energy (Jaggard et al., 2017). Future work should investigate the interplay between sleep plasticity and starvation, and our expression data suggest that sleep deprivation upon starvation in surface fish will lead to increased circulating glucose, while sleep deprivation in response to starvation in cavefish would potentially quickly deplete energy stores.

Sleep restriction in humans is linked to increased food consumption, weight gain and obesity, insulin resistance, and type II diabetes (Cedernaes et al., 2018; Rao et al., 2015; Zhu et al., 2019). Several genes related to glucose homeostasis were upregulated in cave-specific responses. Genes upregulated in Molino cavefish in response to sleep deprivation included: *pyruvate dehydrogenase kinase 4* (*pdk4*), which is increased in mouse models of insulin resistance and type II diabetes (Cedernaes et al., 2018); *carnitine palmitoyltransferase 1b* (*cpt1b*), which is also involved in insulin resistance; and *leptin receptor*, which is involved in a variety of metabolic phenotypes including obesity. Likewise, in Tinaja cavefish genes upregulated in response to sleep deprivation included *insulin receptor substrate 1* (*irs1*) and *solute carrier family 2 member 4* (*slc2a4*) which are both strongly associated with glucose homeostasis. Cavefish populations exhibit intraspecific variation for metabolic phenotypes across independently evolved cave populations (Aspiras et al., 2015; Riddle et al., 2018), and the diverse expression changes presented here suggest intraspecific variation in maintaining glucose homeostasis in response to sleep deprivation.

Despite known links between sleep homeostasis and circadian regulation on a molecular and genetic level, many circadian and sleep-associated genes are not differentially expressed between fish in the sleep deprivation and control treatments. Little is known about genetic regulation of sleep homeostasis in *A. mexicanus*, but previous work has documented a number of factors that regulate circadian cycling of certain transcripts (Beale et al., 2013) and sleep regulation (Jaggard et al., 2018). For example, Pachón cavefish evolved enhanced levels of the wake-promoting *hypocretin neuropeptide precursor HCRT*, conferring sleep loss in this population (Jaggard et al., 2018). At least in part, this phenotype is developmentally regulated as elevated homeobox transcription factor LHX9 specifies a greater number of HCRT-neurons in cavefish (Alie et al., 2018). While *HCRT* was not expressed at sufficient levels to be analyzed by our differential expression analysis, *lhx9* was not differentially expressed after sleep deprivation across any of the populations examined, supporting the notion that different molecular processes regulate sleep duration under standard conditions and sleep homeostasis. In addition, a number of genes previously identified as regulating sleep in zebrafish and mammals are also not differentially expressed for any tetra population in response to sleep deprivation, despite being conserved signaling molecules in sleep regulation across the animal phylogeny. These include *aanat2*, an important enzyme for the production of melatonin, gaba receptors (*gabarapa, gabarapl2*), brain derived neurotrophic factor (*bdnf*), adenosine receptors (*adora1a, adora1b, adora2a, adora2b*), adenosine deaminase (*ada, adar*), adenosine kinase (*adka, adkb*) (Holst & Landolt, 2015), NMDA receptors (e.g., *grin* paralogs, *nsmfa*) (Liu, Liu, Tabuchi, & Wu, 2016), *flotillin (flot1*) (Mackiewicz et al., 2007) and dopamine receptors and transporters (*drd1b, drd4a, drd4b, slc6a3*). Likewise, key clock genes (*clockb, per1a, per1b, per2, per3, arntl1a, arntl1b, cry1aa, cry1ab, cry1ba, cry1bb, cry4, roraa, rorab, rorc*), which are often impacted in sleep deprivation studies (Allada et al., 2017; Archer et al., 2014; Archer & Oster, 2015; Borbély et al., 2016; Franken, 2013; Möller-Levet et al., 2013; Uyhelji et al., 2018), are all not differentially expressed for any tetra population. Together, this suggests that genes regulating sleep duration and circadian function under standard conditions are largely unaffected by a single night of sleep deprivation in tetras.

Several considerations must be taken into account in evaluating our study. First, our study examined the effect of a single night of sleep deprivation in fish housed on a standard 14:10 light cycle. While the results suggest relatively limited changes in the number of differentially expressed genes, it is consistent with studies examining the effects of acute sleep deprivation in other animals. For example, a similar number of genes had altered expression levels in humans after various levels of sleep deprivation (Aho et al., 2013; Cedernaes et al., 2018; Pellegrino et al., 2012). We predict that longer-term sleep deprivation for days or chronic insufficient sleep over a number of days may result in more robust changes in gene expression, however, these protocols would also be likely to induced generalized stress (Pellegrino et al., 2012).

Second, our study employed whole-body sampling for RNA-seq from mRNA transcripts. Tissue specific differences are documented to result from sleep deprivation (Cedernaes et al., 2018) and may obscure signal from specific genes (Diessler et al., 2018). For example, *per2* expression increases in sleep deprived mice and remains elevated for varied amounts of time depending on the tissue (Curie, Maret, Emmenegger, & Franken, 2015). At 30dpf, brain dissection is technically challenging and would likely require pooling across samples. Further, precise dissection of tissue takes time and would result in the samples collected last being sleep deprived for longer than the samples collected first.

To our knowledge, these findings are the first genome-wide analysis of sleep deprivation induced changes in fish. In zebrafish, sleep deprivation robustly impacts cellular processes and behavior (Aho et al., 2017; Elbaz et al., 2017; Pinheiro-da-Silva, Silva, Nogueira, & Luchiari, 2017; Zada et al., 2019), but the effects on large-scale changes in gene expression have not been investigated. Both zebrafish and *A. mexicanus* provide robust models for the identification of genetic and pharmacological regulators of sleep (Duboué et al., 2012; Jaggard et al., 2018; Prober, 2018; Rihel, Prober, & Schier, 2010), suggesting these models can be used to investigate the genetic architecture associated with sleep loss. Our findings, that sleep deprivation induces different molecular signatures in each of the four *A. mexicanus* populations tested, raises the possibility that the response to sleep pressure is highly heterogenous across individuals of the same species. The application of recently developed gene-editing approaches in *A. mexicanus* (Ma et al., 2018; Stahl et al., 2019) combined with the behavioral assay described here may allow for functional validation of these genes and identification of novel regulators of sleep homeostasis.

## Acknowledgements

We thank the University of Minnesota Genomics Center for their guidance and performing the cDNA library preparations and Illumina HiSeq 2500 sequencing. The Minnesota Supercomputing Institute (MSI) at the University of Minnesota provided resources that contributed to the research results reported within this paper. Funding was supported by NIH (1R01GM127872-01 to SEM and ACK) and NSF award IOS 165674 to ACK, and a US-Israel BSF award to ACK. CNP was supported by Grand Challenges in Biology Postdoctoral Program at the University of Minnesota College of Biological Sciences. Institutional Animal Care and Use Committee at Florida Atlantic University (Protocol #A15-32).

## Data availability statement

All RNAseq reads are available on the SRA (Accession numbers given in Table S1). Raw expression counts data are given as a supplementary file.

## References

Aho, V., Ollila, H. M., Rantanen, V., Kronholm, E., Surakka, I., van Leeuwen, W. M., … Härmä, M. (2013). Partial sleep restriction activates immune response-related gene expression pathways: experimental and epidemiological studies in humans. PloS one, 8(10), e77184.

Aho, V., Vainikka, M., Puttonen, H. A., Ikonen, H. M., Salminen, T., Panula, P., … Wigren, H. K. (2017). Homeostatic response to sleep/rest deprivation by constant water flow in larval zebrafish in both dark and light conditions. Journal of sleep research, 26(3), 394–400.

Alie, A., Devos, L., Torres-Paz, J., Prunier, L., Boulet, F., Blin, M., … Retaux, S. (2018). Developmental evolution of the forebrain in cavefish, from natural variations in neuropeptides to behavior. eLife, 7, e32808.

Allada, R., Cirelli, C., & Sehgal, A. (2017). Molecular mechanisms of sleep homeostasis in flies and mammals. Cold Spring Harbor perspectives in biology, 9(8), a027730.

Anafi, R. C., Kayser, M. S., & Raizen, D. M. (2018). Exploring phylogeny to find the function of sleep. Nature Reviews Neuroscience, 1.

Andrews, S. (2014). FastQC: a quality control tool for high throughput sequence data. Version 0.11. 2. Babraham Institute, Cambridge, UK http://www.bioinformatics.babraham.ac.uk/projects/fastqc.

Archer, S. N., Laing, E. E., Moller-Levet, C. S., van der Veen, D. R., Bucca, G., Lazar, A. S., … Dijk, D. J. (2014). Mistimed sleep disrupts circadian regulation of the human transcriptome. Proceedings of the National Academy of Sciences USA, 111(6), E682–691. doi:10.1073/pnas.1316335111

Archer, S. N., & Oster, H. (2015). How sleep and wakefulness influence circadian rhythmicity: effects of insufficient and mistimed sleep on the animal and human transcriptome. Journal of sleep research, 24(5), 476–493. doi:10.1111/jsr.12307

Aspiras, A. C., Rohner, N., Martineau, B., Borowsky, R. L., & Tabin, C. J. (2015). *Melanocortin 4 receptor* mutations contribute to the adaptation of cavefish to nutrient-poor conditions. Proceedings of the National Academy of Sciences, 112(31), 9668–9673.

Beale, A., Guibal, C., Tamai, T. K., Klotz, L., Cowen, S., Peyric, E., … Whitmore, D. (2013). Circadian rhythms in Mexican blind cavefish *Astyanax mexicanus* in the lab and in the field. Nature communications, 4, 2769–2769. doi:10.1038/ncomms3769

Blask, D. E. (2009). Melatonin, sleep disturbance and cancer risk. Sleep medicine reviews, 13(4), 257–264.

Bolger, A. M., Lohse, M., & Usadel, B. (2014). Trimmomatic: A flexible trimmer for Illumina sequence data. Bioinformatics, 30(15), 2114–2120. doi:10.1093/bioinformatics/btu170

Borbély, A. A., Daan, S., Wirz-Justice, A., & Deboer, T. (2016). The two-process model of sleep regulation: a reappraisal. Journal of sleep research, 25(2), 131–143.

Borowsky, R. (2008). *Astyanax mexicanus*, the blind Mexican cave fish: A model for studies in development and morphology. Cold Spring Harbor Protocols, 2008(11), pdb. emo107.

Bradic, M., Beerli, P., Garcia-de Leon, F. J., Esquivel-Bobadilla, S., & Borowsky, R. L. (2012). Gene flow and population structure in the Mexican blind cavefish complex *(Astyanax mexicanus)*. BMC evolutionary biology, 12(1), 9.

Cappuccio, F. P., D’elia, L., Strazzullo, P., & Miller, M. A. (2010). Quantity and quality of sleep and incidence of type 2 diabetes: a systematic review and meta-analysis. Diabetes care, 33(2), 414–420.

Cedernaes, J., Schönke, M., Westholm, J. O., Mi, J., Chibalin, A., Voisin, S.,Dickson, S. L. (2018). Acute sleep loss results in tissue-specific alterations in genome-wide DNA methylation state and metabolic fuel utilization in humans. Science advances, 4(8), eaar8590.

Cirelli, C., Faraguna, U., & Tononi, G. (2006). Changes in brain gene expression after long-term sleep deprivation. Journal of neurochemistry, 98(5), 1632–1645.

Coghill, L. M., Hulsey, C. D., Chaves-Campos, J., García de Leon, F. J., & Johnson, S. G. (2014). Next generation phylogeography of cave and surface *Astyanax mexicanus*. Molecular Phylogenetics and Evolution, 79, 368–374.

Conde, J., Otero, M., Scotece, M., Abella, V., López, V., Pino, J., … Gualillo, O. (2016). E74-like factor 3 and nuclear factor-κB regulate lipocalin-2 expression in chondrocytes. The Journal of physiology, 594(21), 6133–6146.

Cruickshank, T. E., & Hahn, M. W. (2014). Reanalysis suggests that genomic islands of speciation are due to reduced diversity, not reduced gene flow. Molecular ecology, 23, 3133–3157.

Curie, T., Maret, S., Emmenegger, Y., & Franken, P. (2015). In vivo imaging of the central and peripheral effects of sleep deprivation and suprachiasmatic nuclei lesion on PERIOD-2 protein in mice. Sleep, 38(9), 1381–1394.

Davies, S. K., Ang, J. E., Revell, V. L., Holmes, B., Mann, A., Robertson, F. P., … Skene, D. J. (2014). Effect of sleep deprivation on the human metabolome. Proceedings of the National Academy of Sciences USA, 111(29), 10761–10766. doi:10.1073/pnas.1402663111

Dickinson, M. E., Flenniken, A. M., Ji, X., Teboul, L., Wong, M. D., White, J. K., … Adissu, H. (2016). High-throughput discovery of novel developmental phenotypes. Nature, 537(7621), 508.

Diessler, S., Jan, M., Emmenegger, Y., Guex, N., Middleton, B., Skene, D. J., … Pagni, M. (2018). A systems genetics resource and analysis of sleep regulation in the mouse. PLoS biology, 16(8), e2005750.

Dobin, A., Davis, C. A., Schlesinger, F., Drenkow, J., Zaleski, C., Jha, S., … Gingeras, T. R. (2013). STAR: ultrafast universal RNA-seq aligner. Bioinformatics, 29(1), 15–21.

Duboué, E. R., Borowsky, R. L., & Keene, A. C. (2012). β-adrenergic signaling regulates evolutionarily derived sleep loss in the mexican cavefish. Brain, Behavior, and Evolution, 80, 233–243. doi:10.1159/000341403

Duboué, E. R., Keene, A. C., & Borowsky, R. L. (2011). Evolutionary convergence on sleep loss in cavefish populations. Current Biology, 21(8), 671–676. doi:10.1016/j.cub.2011.03.020

Elbaz, I., Zada, D., Tovin, A., Braun, T., Lerer-Goldshtein, T., Wang, G., … Appelbaum, L. (2017). Sleep-dependent structural synaptic plasticity of inhibitory synapses in the dendrites of hypocretin/orexin neurons. Molecular neurobiology, 54(8), 6581–6597.

Elliott, A. S., Huber, J. D., O’Callaghan, J. P., Rosen, C. L., & Miller, D. B. (2014). A review of sleep deprivation studies evaluating the brain transcriptome. SpringerPlus, 3(1), 728.

Espinasa, L., Rivas-Manzano, P., & Pérez, H. E. (2001). A new blind cave fish population of genus Astyanax: geography, morphology and behavior. Environmental Biology of Fishes, 62(1-3), 339–344.

Fariello, M. I., Boitard, S., Naya, H., SanCristobal, M., & Servin, B. (2013). Detecting signatures of selection through haplotype differentiation among hierarchically structured populations. Genetics, 193(3), 929–941.

Franken, P. (2013). A role for clock genes in sleep homeostasis. Current opinion in neurobiology, 23, 864–872.

Gajulapalli, V. N. R., Samanthapudi, V. S. K., Pulaganti, M., Khumukcham, S. S., Malisetty, V. L., Guruprasad, L., … Manavathi, B. (2016). A transcriptional repressive role for epithelial-specific ETS factor ELF3 on oestrogen receptor alpha in breast cancer cells. Biochemical Journal, 473(8), 1047–1061.

Gross, J. B., Furterer, A., Carlson, B. M., & Stahl, B. A. (2013). An integrated transcriptome-wide analysis of cave and surface dwelling *Astyanax mexicanus*. PloS one, 8(2), e55659.

Han, F., Lin, L., Li, J., Aran, A., Dong, S. X., An, P., … Wang, J. S. (2012). TCRA, P2RY11, and CPT1B/CHKB associations in Chinese narcolepsy. Sleep medicine, 13(3), 269–272.

Haus, E. L., & Smolensky, M. H. (2013). Shift work and cancer risk: potential mechanistic roles of circadian disruption, light at night, and sleep deprivation. Sleep medicine reviews, 17(4), 273–284.

Herman, A., Brandvain, Y., Weagley, J., Jeffery, W. R., Keene, A. C., Kono, T. J. Y., … McGaugh, S. E. (2018). The role of gene flow in rapid and repeated evolution of cave related traits in Mexican tetra, *Astyanax mexicanus*. Molecular ecology, 27, 4397–4416

Hinaux, H., Poulain, J., Da Silva, C., Noirot, C., Jeffery, W. R., Casane, D., & Retaux, S. (2013). De novo sequencing of *Astyanax mexicanus* surface fish and Pachón cavefish transcriptomes reveals enrichment of mutations in cavefish putative eye genes. PloS one, 8, e53553.

Holst, S. C., & Landolt, H.-P. (2015). Sleep homeostasis, metabolism, and adenosine. Current Sleep Medicine Reports, 1(1), 27–37.

Homan, E. A., Kim, Y.-G., Cardia, J. P., & Saghatelian, A. (2011). Monoalkylglycerol ether lipids promote adipogenesis. Journal of the American Chemical Society, 133(14), 5178–5181.

Igarashi, M., Osuga, J.-i., Uozaki, H., Sekiya, M., Nagashima, S., Takahashi, M., … Ohta, K. (2010). The critical role of neutral cholesterol ester hydrolase 1 in cholesterol removal from human macrophages. Circulation research, 107(11), 1387–1395.

Jaggard, J. B., Lloyd, E., Lopatto, A., Duboue, E. R., & Keene, A. C. (2019). Automated measurements of sleep and locomotor activity in Mexican cavefish. JoVE (Journal of Visualized Experiments)(145), e59198.

Jaggard, J. B., Robinson, B. G., Stahl, B. A., Oh, I., Masek, P., Yoshizawa, M., & Keene, A. C. (2017). The lateral line confers evolutionarily derived sleep loss in the Mexican cavefish. Journal of Experimental Biology, 220, 284–293.

Jaggard, J. B., Stahl, B. A., Lloyd, E., Prober, D. A., Duboue, E. R., & Keene, A. C. (2018). Hypocretin underlies the evolution of sleep loss in the Mexican cavefish. eLife, 7, e32637.

Joiner, W. J. (2016). Unraveling the evolutionary determinants of sleep. Current Biology, 26(20), R1073–R1087.

Keene, A., Yoshizawa, M., & McGaugh, S. E. (2015). Biology and Evolution of the Mexican Cavefish: Elsevier: Academic Press.

Keene, A. C., & Duboue, E. R. (2018). The origins and evolution of sleep. Journal of Experimental Biology, 221(11), jeb159533.

Kim, J., Okamoto, H., Huang, Z., Anguiano, G., Chen, S., Liu, Q., … Hamid, R. (2017). Amino acid transporter Slc38a5 controls glucagon receptor inhibition-induced pancreatic α cell hyperplasia in mice. Cell metabolism, 25(6), 1348–1361. e1348.

Kwon, M.-c., Koo, B.-K., Kim, Y.-Y., Lee, S.-H., Kim, N.-S., Kim, J.-H., & Kong, Y.-Y. (2009). Essential role of CR6-interacting factor 1 (Crif1) in E74-like factor 3 (ELF3)-mediated intestinal development. Journal of Biological Chemistry, 284(48), 33634–33641.

Lane, J. M., Liang, J., Vlasac, I., Anderson, S. G., Bechtold, D. A., Bowden, J., … Luik, A. I. (2017). Genome-wide association analyses of sleep disturbance traits identify new loci and highlight shared genetics with neuropsychiatric and metabolic traits. Nature genetics, 49(2), 274.

Leproult, R., Holmbäck, U., & Van Cauter, E. (2014). Circadian misalignment augments markers of insulin resistance and inflammation, independently of sleep loss. Diabetes, DB_131546.

Levy, P., Tamisier, R., Arnaud, C., Monneret, D., Baguet, J., Stanke-Labesque, F., … Pepin, J. (2012). Sleep deprivation, sleep apnea and cardiovascular diseases. Frontiers in bioscience (Elite edition), 4, 2007–2021.

Libourel, P.-A., Barrillot, B., Arthaud, S., Massot, B., Morel, A.-L., Beuf, O., … Luppi, P.-H. (2018). Partial homologies between sleep states in lizards, mammals, and birds suggest a complex evolution of sleep states in amniotes. PLoS biology, 16(10), e2005982.

Liu, S., Croniger, C., Arizmendi, C., Harada-Shiba, M., Ren, J., Poli, V., … Friedman, J. E. (1999). Hypoglycemia and impaired hepatic glucose production in mice with a deletion of the C/EBPβ gene. The journal of clinical Investigation, 103(2), 207–213.

Liu, S., Liu, Q., Tabuchi, M., & Wu, M. N. (2016). Sleep drive is encoded by neural plastic changes in a dedicated circuit. Cell, 165(6), 1347–1360.

Ma, L., Strickler, A. G., Parkhurst, A., Yoshizawa, M., Shi, J., & Jeffery, W. R. (2018). Maternal genetic effects in *Astyanax* cavefish development. Developmental Biology, 441(2), 209–220.

Mackiewicz, M., Shockley, K. R., Romer, M. A., Galante, R. J., Zimmerman, J. E., Naidoo, N., … Pack, A. I. (2007). Macromolecule biosynthesis: a key function of sleep. Physiological Genomics, 31(3), 441–457.

Mackiewicz, M., Zimmerman, J. E., Shockley, K. R., Churchill, G. A., & Pack, A. I. (2009). What are microarrays teaching us about sleep? Trends in Molecular Medicine, 15(2), 79–87. doi:10.1016/j.molmed.2008.12.002

McGaugh, S. E., Gross, J. B., Aken, B., Blin, M., Borowsky, R., Chalopin, D., … Warren, W. C. (2014). The cavefish genome reveals candidate genes for eye loss. Nature communications, 5, 5307–5307.

Miyagawa, T., Kawashima, M., Nishida, N., Ohashi, J., Kimura, R., Fujimoto, A., … Lin, L. (2008). Variant between CPT1B and CHKB associated with susceptibility to narcolepsy. Nature genetics, 40(11), 1324.

Möller-Levet, C. S., Archer, S. N., Bucca, G., Laing, E. E., Slak, A., Kabiljo, R., … Smith, C. P. (2013). Effects of insufficient sleep on circadian rhythmicity and expression amplitude of the human blood transcriptome. Proceedings of the National Academy of Sciences, 110(12), E1132-E1141.

Nakazaki, C., Noda, A., Koike, Y., Yamada, S., Murohara, T., & Ozaki, N. (2012). Association of insomnia and short sleep duration with atherosclerosis risk in the elderly. American journal of hypertension, 25(11), 1149–1155.

Nath, R. D., Bedbrook, C. N., Abrams, M. J., Basinger, T., Bois, J. S., Prober, D. A., … Goentoro, L. (2017). The jellyfish *Cassiopea* exhibits a sleep-like state. Current Biology, 27(19), 2984–2990. e2983.

Obholzer, N., Wolfson, S., Trapani, J. G., Mo, W., Nechiporuk, A., Busch-Nentwich, E., … Duncan, R. N. (2008). Vesicular glutamate transporter 3 is required for synaptic transmission in zebrafish hair cells. Journal of Neuroscience, 28(9), 2110–2118.

Okazaki, H., Igarashi, M., Nishi, M., Sekiya, M., Tajima, M., Takase, S., … Okazaki, S. (2008). Identification of neutral cholesterol ester hydrolase, a key enzyme removing cholesterol from macrophages. Journal of Biological Chemistry, 283(48), 33357–33364.

Passow, C. N., Kono, T. J., Stahl, B. A., Jaggard, J. B., Keene, A. C., & McGaugh, S. E. (2019). Nonrandom RNAseq gene expression associated with RNAlater and flash freezing storage methods. Molecular Ecology Resources, 19(2), 456–464.

Pellegrino, R., Sunaga, D. Y., Guindalini, C., Martins, R. C., Mazzotti, D. R., Wei, Z., … Tufik, S. (2012). Whole blood genome-wide gene expression profile in males after prolonged wakefulness and sleep recovery. Physiological Genomics, 44(21), 1003–1012.

Pertea, M., Kim, D., Pertea, G. M., Leek, J. T., & Salzberg, S. L. (2016). Transcript-level expression analysis of RNA-seq experiments with HISAT, StringTie and Ballgown. Nature protocols, 11(9), 1650.

Pertea, M., Pertea, G. M., Antonescu, C. M., Chang, T.-C., Mendell, J. T., & Salzberg, S. L. (2015). StringTie enables improved reconstruction of a transcriptome from RNA-seq reads. Nature biotechnology, 33(3), 290.

Pinheiro-da-Silva, J., Silva, P. F., Nogueira, M. B., & Luchiari, A. C. (2017). Sleep deprivation effects on object discrimination task in zebrafish *(Danio rerio)*. Animal cognition, 20(2), 159–169.

Podobed, P. S., Alibhai, F. J., Chow, C.-W., & Martino, T. A. (2014). Circadian regulation of myocardial sarcomeric Titin-cap (Tcap, telethonin): identification of cardiac clock-controlled genes using open access bioinformatics data. PloS one, 9(8), e104907.

Prober, D. A. (2018). Discovery of hypocretin/orexin ushers in a new era of sleep research. Trends in neurosciences, 41(2), 70–72.

Rao, M. N., Neylan, T. C., Grunfeld, C., Mulligan, K., Schambelan, M., & Schwarz, J.-M. (2015). Subchronic sleep restriction causes tissue-specific insulin resistance. The Journal of Clinical Endocrinology & Metabolism, 100(4), 1664–1671.

Riddle, M. R., Aspiras, A. C., Gaudenz, K., Peuß, R., Sung, J. Y., Martineau, B., … McGaugh, S. (2018). Insulin resistance in cavefish as an adaptation to a nutrient-limited environment. Nature, 555(7698), 647.

Rihel, J., Prober, D. A., & Schier, A. F. (2010). Monitoring sleep and arousal in zebrafish. In Methods in Cell Biology (Vol. 100, pp. 281–294): Elsevier.

Robinson, M. D., McCarthy, D. J., & Smyth, G. K. (2010). edgeR: a Bioconductor package for differential expression analysis of digital gene expression data. Bioinformatics, 26(1), 139–140.

Schlamp, F., Van Der Made, J., Stambler, R., Chesebrough, L., Boyko, A. R., & Messer, P. W. (2016). Evaluating the performance of selection scans to detect selective sweeps in domestic dogs. Molecular ecology, 25(1), 342–356. doi:10.1111/mec.13485

Shatnawi, A., Norris, J., Chaveroux, C., Jasper, J., Sherk, A., McDonnell, D., & Giguere, V. (2014). ELF3 is a repressor of androgen receptor action in prostate cancer cells. Oncogene, 33(7), 862.

Stahl, B. A., & Gross, J. B. (2017). A comparative transcriptomic analysis of development in two *Astyanax* cavefish populations. Journal of Experimental Zoology Part B: Molecular and Developmental Evolution, 328, 515–532.

Stahl, B. A., Peuß, R., McDole, B., Kenzior, A., Jaggard, J. B., Gaudenz, K., … Keene, A. C. (2019). Stable transgenesis in *Astyanax mexicanus* using the Tol2 transposase system. Developmental Dynamics.

Toyama, R., Chen, X., Jhawar, N., Aamar, E., Epstein, J., Reany, N., … Dawid, I. B. (2009). Transcriptome analysis of the zebrafish pineal gland. Developmental dynamics: an official publication of the American Association of Anatomists, 238(7), 1813–1826.

Uyhelji, H. A., Kupfer, D. M., White, V. L., Jackson, M. L., Van Dongen, H. P., & Burian, D. M. (2018). Exploring gene expression biomarker candidates for neurobehavioral impairment from total sleep deprivation. BMC genomics, 19(1), 341.

Wang, J., Chakrabarty, S., Bui, Q., Ruf, W., & Samad, F. (2015). Hematopoietic tissue factor–protease-activated receptor 2 signaling promotes hepatic inflammation and contributes to pathways of gluconeogenesis and steatosis in obese mice. The American journal of pathology, 185(2), 524–535.

Yachida, S., Wood, L. D., Suzuki, M., Takai, E., Totoki, Y., Kato, M., … Hama, N. (2016). Genomic sequencing identifies ELF3 as a driver of ampullary carcinoma. Cancer cell, 29(2), 229–240.

Yeung, T.-L., Leung, C. S., Wong, K.-K., Gutierrez-Hartmann, A., Kwong, J., Gershenson, D. M., & Mok, S. C. (2017). ELF3 is a negative regulator of epithelial-mesenchymal transition in ovarian cancer cells. Oncotarget, 8(10), 16951.

Yoshizawa, M. (2015). Behaviors of cavefish offer insight into developmental evolution. Molecular Reproduction and Development, 82(4), 268–280. doi:10.1002/mrd.22471

Yoshizawa, M., Robinson, B., Duboue, E. R., Masek, P., Jaggard, J. B., O’Quin, K. E., … Keene, A. C. (2015). Distinct genetic architecture underlies the emergence of sleep loss and prey-seeking behavior in the Mexican cavefish. BMC Biology, 13(1), 1.

Yoshizawa, M., Settle, A., Hermosura, M. C., Tuttle, L. J., Cetraro, N., Passow, C. N., & McGaugh, S. E. (2018). The evolution of a series of behavioral traits is associated with autism-risk genes in cavefish. BMC evolutionary biology, 18(1), 89.

Zada, D., Bronshtein, I., Lerer-Goldshtein, T., Garini, Y., &; Appelbaum, L. (2019). Sleep increases chromosome dynamics to enable reduction of accumulating DNA damage in single neurons. Nature communications, 10(1), 895.

Zhang, C., Kho, Y.-S., Wang, Z., Chiang, Y. T., Ng, G. K., Shaw, P.-C Qi, R. Z. (2014). Transmembrane and coiled-coil domain family 1 is a novel protein of the endoplasmic reticulum. PloS one, 9(1), e85206.

Zhu, B., Shi, C., Park, C. G., Zhao, X., &; Reutrakul, S. (2019). Effects of sleep restriction on metabolism-related parameters in healthy adults: A comprehensive review and meta-analysis of randomized controlled trials. Sleep medicine reviews, 45, 18–30.

